# Strategies for understanding dynamic, personalized profiles of host-derived proteins and microbes from human stool

**DOI:** 10.1101/551143

**Authors:** Ellen Casavant, Les Dethlefsen, Kris Sankaran, Daniel Sprockett, Susan Holmes, David A. Relman, Joshua E. Elias

## Abstract

Measuring host proteins through noninvasive stool-based assays opens new avenues for characterizing states of gastrointestinal health. However, the extent to which these proteins vary over time and between healthy subjects is poorly characterized. Here, we characterize technical and biological sources of variability in mass spectrometry-based measurements of host proteins in stool. We identify the proteins that most vary over time within an individual, and among different individuals. Finally, we examine and compare temporal and inter-individual variation in host protein and bacterial taxonomic profiles of the same fecal specimens. To address these issues, five self-reported healthy individuals were each sampled eight times over four weeks. First, we demonstrate that mass spectrometry-based identification and label-free quantification of stool proteins exhibit non-significant variability (p>0.05) between both technical and preparative replicates for a subset of 78 proteins, supporting the utility of this method for biomarker measurement. Second, although 13 human stool proteins varied significantly in relative abundance over time within individuals, 58 proteins varied significantly (at least four-fold) between subjects. The average pair-wise difference between individuals was greater than the average within-subject difference for both the proteome and microbiome datasets (p<0.0001). Fecal host proteins, like the traditional fecal protein marker, calprotectin, unambiguously pointed to innate and adaptive immune responses. For example, one subject’s fecal protein profile suggested a sub-clinical inflammatory state. From these data, we conclude that host-centric protein measurements in stool reveal a wide range of variation during states of apparent health, and add a valuable complementary insight into host-microbiota relationships.

**IMPORTANCE:** Human proteins in stool hold untapped potential for characterizing gastrointestinal health. To fully harness this potential and create a baseline of healthy stool protein abundances and identifications, it will be important to establish the extent to which these proteins might vary in the absence of disease. This study quantifies the major sources of variation in stool protein abundance data. We assessed technical, preparative, temporal, and inter-subject variability of human protein abundances in stool and found that among these sources, differences between subjects accounted for the greatest amount of variation, followed by temporal differences, and then technical factors. Our paired microbiome analysis found matching patterns of temporal and inter-subject variability. By characterizing multiple variance parameters in host stool protein abundances, our analysis helps to contextualize a wide range of future disease-focused stool studies as well as elucidate host-microbe interactions.

## 1. Introduction

The expression of host proteins in the human gastrointestinal tract contributes to the health of that ecosystem, helping to regulate the diverse communities of microbes whose composition and activities in turn affect host physiology. By understanding better the identity of these proteins and their relative abundances we gain another window, besides that afforded by assessments of the gut microbiota, into the mechanisms that underlie important homeostatic processes in the gastrointestinal tract. Stool is the most easily obtained specimen type from this organ. Yet, variation in the abundances of the hundreds of host proteins found in human stool is poorly understood, in part because their relative abundances in stool reflect the contributions of multiple host cell types throughout the gastrointestinal tract, different mechanisms of secretion and release—including sloughing of epithelium, different protein half-lives, diverse processes of protein degradation, and varying gut transit times. But in addition, there are surprisingly few, broad surveys of host proteins in human stool during states of health. We reasoned that the characterization of human proteins in stool collected from healthy donors has important implications for precision health. Thus, this non-invasive approach may yield new opportunities for improved diagnosis and recognition of pre-clinical gastrointestinal health states.

Prior efforts to understand gastrointestinal health have focused on the gut microbiota. Trillions of microbes living inside the human gastrointestinal tract exert a strong influence on host health, both locally in the intestinal tract, and throughout the body. Microbial community surveys using 16S rRNA sequence data can track perturbations such as antibiotic exposure and reveal connections between the composition of the microbiome and health conditions such as obesity, diabetes, and colorectal cancer (1–7). While next-generation sequencing technologies have shed new light on how microbial communities vary over time and across health states, the effects of particular microbial species on human health are less clear. Likewise, the dynamic responses of gastrointestinal cells and tissues to changes in the resident microbial communities are poorly understood. Thus, targeted study of extracellular host proteins recovered from stool (henceforth referred to as stool proteins) stands to add an often-overlooked dimension to the characterization of host-microbe interactions.

Our prior studies in laboratory mice have shown that distinct sets of stool proteins are associated with different microbial communities (8) and with different states of inflammation (9). In addition, we have shown that host stool proteins could distinguish different healthy human donors (8) – a finding that was subsequently extended in a comparison of obese and lean clinical study participants (10). To further characterize the utility of human stool proteins, we addressed in the present study, the degree of variance in human stool protein abundances among multiple donors over multiple time scales.

We argue that establishing the parameters of a healthy gastrointestinal tract proteome is an important prerequisite for identifying the stool proteins and their abundances that might be capable of defining a wide range of health. Since human stool specimens are inherently generated under uncontrolled, non-laboratory conditions, the ability to extrapolate from studies of protein dynamics in laboratory animals is fundamentally limited. We expect that the confluence of diet, exercise, and environmental stressors cause stool protein abundances to vary significantly over time and between individuals. This should affect their utility as reliable biomedical diagnostic tools.

Here, we compare stool proteins and microbial community compositions from five subjects surveyed over eight time points spanning four weeks. We developed a statistical approach that contends with the stochasticity inherent to these kinds of shotgun proteomics studies, and that enabled three key findings. First, we found that technical sources of measurement variation were largely insignificant (log(p) > −1.30), an essential prerequisite for performing large-scale, clinical studies. Second, we confirm that host-expressed stool proteins vary over time, but that variation between subjects is far greater than this intra-host temporal variability. Third, by comparing protein profiles to parallel microbial 16S rRNA gene sequence surveys, we found that while they reveal comparable subject-specific profiles, the proteomic data provide complementary insight into human inflammation.

## 2. Results

### 2.1 Parallel stool protein and microbe surveys for assessing sources of measurement variability

We designed a human stool sampling protocol that allowed us to assess measurement variation over time and between subjects. We surveyed stool from five self-reported healthy subjects over four weeks. Each subject provided one stool specimen on the same two consecutive days of each week for four weeks, totaling eight specimens per subject (**Fig. 1A**). To distinguish biological sources of stool protein variation from technical sources, we prepared each specimen in duplicate (preparative replicates) and subjected each preparative replicate to two replicate LC-MS/MS analyses (technical replicates) (**Fig. 1B**). A third preparative replicate was obtained for microbiome sequencing. All subject, day, and preparative replicates were analyzed by LC-MS/MS in a randomized fashion (**Materials and Methods**). Non-significant variation of technical and preparative variation (log(p) > −1.30) between proteins in experiments as described below supports the validity of these stool protein measurements. Microbial temporal and inter-individual trends were measured in parallel using 16S rRNA V4-V5 amplicon surveys without replication.

**Figure 1:**
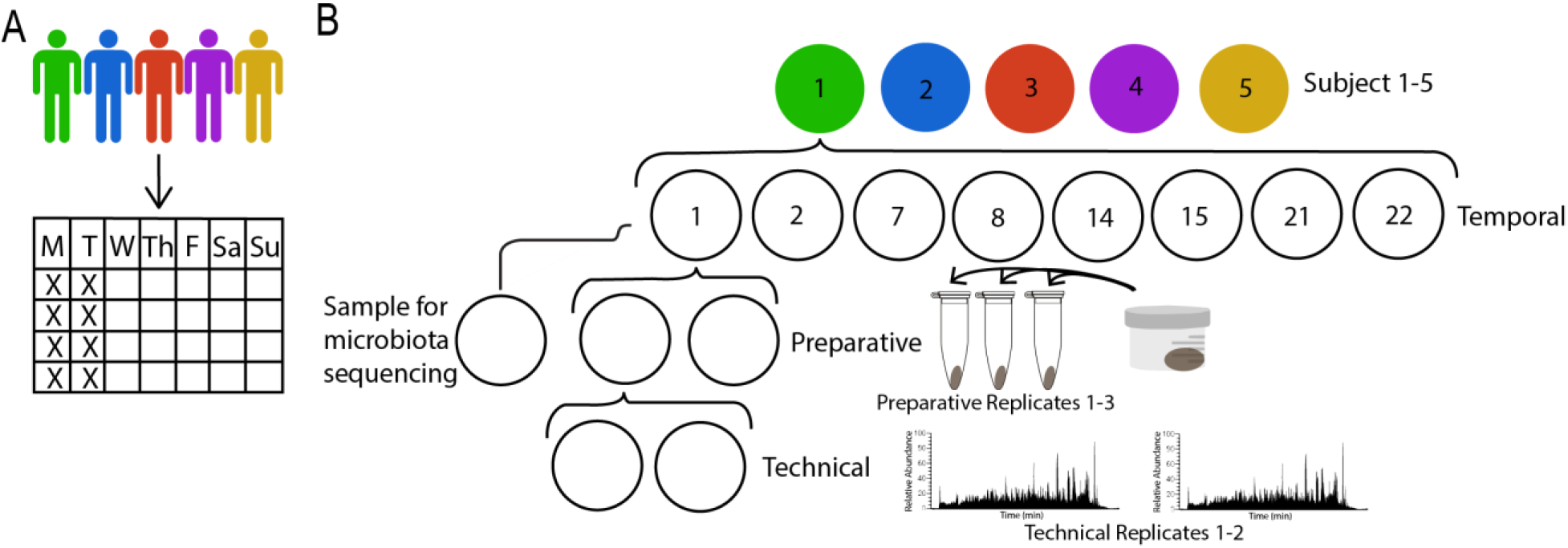
Stool sampling strategy. **A**. Five self-reported healthy subjects provided two stool specimens on two consecutive days of four consecutive weeks. **B**. Each stool specimen was aliquoted and prepared in duplicate (preparative replicates). Each preparative replicate from all 40 specimens were processed in a random order and subjected to two LC-MS analyses (technical replicates). A third stool specimen aliquot underwent 16srRNA gene sequencing for microbe enumeration.

We measured 456 stool proteins across all subjects and time points (protein FDR <1.5%; **Fig. 2A**). Total unique protein identifications per subject ranged from 256 to 384. Several prominent protein functional groups were enriched above the expected presence in the entire human proteome among these proteins, including immunoglobulins (4.3-fold), serine proteases (7.4-fold), and protease inhibitors (9.76-fold)(**Fig. 2B**). These protein classes have clear gastrointestinal relevance, and support our method’s ability to enrich for proteins mediating interactions between the host, gut microbes, and diet (8).

**Figure 2:**
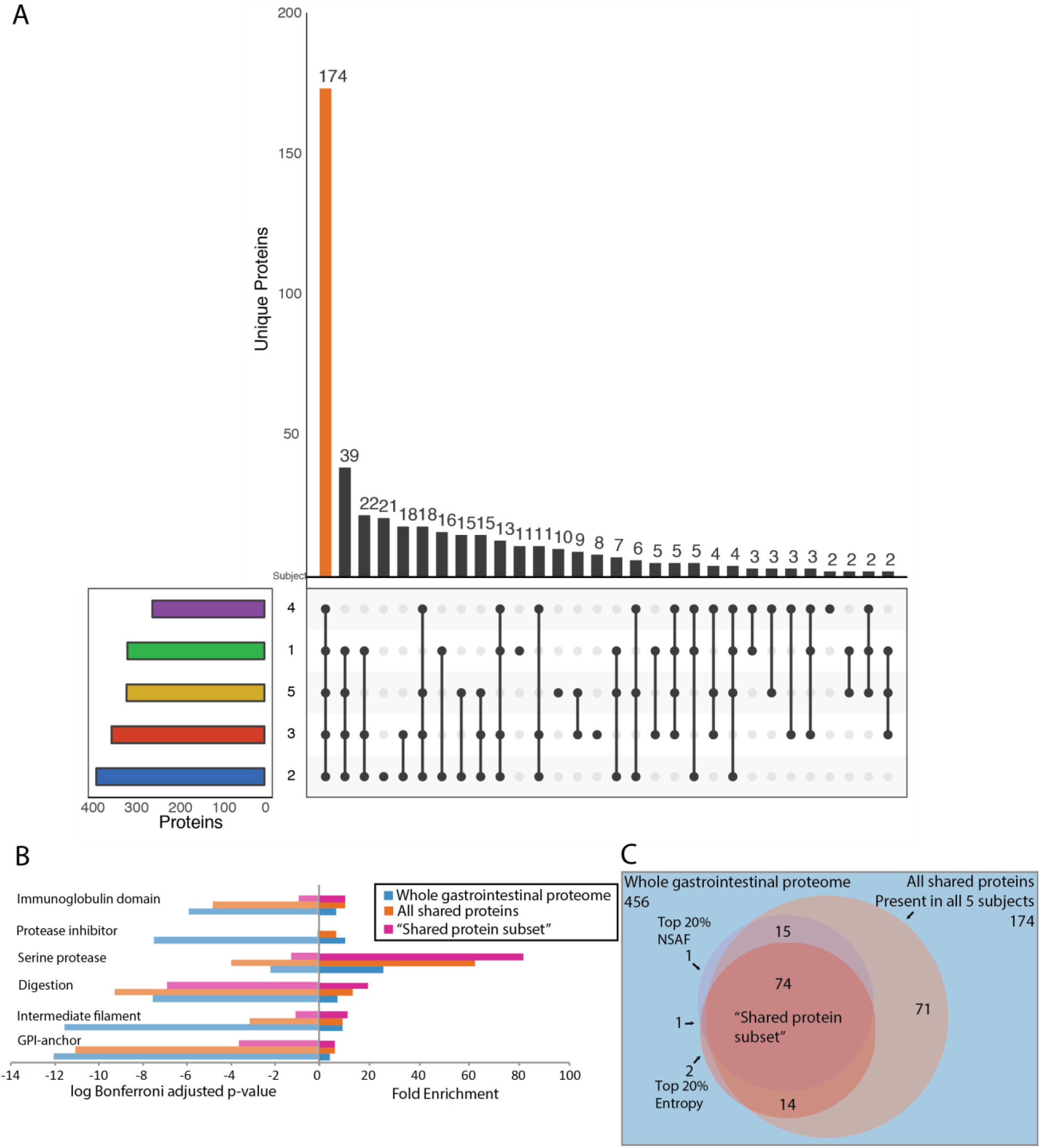
Towards a representative shared protein subset of extracellular host-derived stool proteins. **A**. Most protein identifications were specific to each donor. The subset plot (“UpSetR” version 1.3.3) shows the proteins shared between each subject. 174 proteins were identified from at least one time point across all subjects (orange bar). 11% of the data set (52 proteins) were identified in just one subject. **B**. The shared protein subset is the union between the 174 shared proteins, the top 20% summed (across all specimen in the dataset) NSAF proteins, and the top 20% summed (across all specimen in the dataset) entropy proteins. This union totals 74 proteins, which compose the shared protein subset used in subsequent ANOVA and logistic regression tests (**Figure 3**). **C**. Proteolysis and immune-related proteins were significantly enriched in three data subsets: the entire extracellular host stool protein data set generated here, the 180 protein shared between all donors, and the 74 shared protein subset proteins. Digestive and serine protease proteins showed the greatest enrichment among the shared protein subset proteins. Intracellular protein groups like GI-anchor and intermediate filaments showed no representation among shared protein subset proteins. Gene ontology analysis was conducted using the DAVID web portal (29). Noted protein subsets were input as a “foreground” gene list and compared to a background genelist consisting of the entire default human proteome. The resulting gene ontologies exceeding a Bonferroni-corrected p-value of 0.05 were considered significantly enriched; a subset of which are shown here.

Our parallel 16S rRNA survey enumerated 1,770 microbial amplicon sequence variants (ASVs) across all subjects and time points, with 320 to 808 unique ASVs identified per subject (**Supp. Fig. 1**). ASVs were associated with nine bacterial and one archaeal phyla, with Firmicutes (range 54%-79%, mean 70%) and Bacteroidetes (range 6%-31%, mean 18%) dominating the gut microbiota of all subjects (**Supp. Fig. 2**). Actinobacteria (0.5%-3.5%), Proteobacteria (0.4%-1.2%) and Verrucomicrobia (0.05%-28%) were also detected in all subjects. Euryarchaeota, the archaeal phylum containing methanogens, was detected in two subjects (3.0% and 7.1%).

Phylum-level relative abundances have been reported to vary widely and are sensitive to cell lysis procedures (11). Nevertheless, our results are consistent with the ranges reported previously for healthy adults (12, 13). At a finer scale, both the microbial families and species we detected, and their variability over time within subjects, are similar to published results from healthy adults (**Supp Fig 3**) (14, 15). With confidence in both protein and microbe datasets, we chose to first examine all types of stool protein variability, before comparing patterns with our microbe dataset.

### 2.2 A shared protein subset for testing multiple sources of variability

The gastrointestinal and immunological functions that were enriched among the stool proteins we measured suggested their diagnostic utility for characterizing a range of health states. However, for our proteomic approach to have clinical utility, we first sought to establish confidence parameters for the technical and biological variability in their measurement. To accomplish this, we tested how consistently proteins were observed and, when observed, how consistent their measured abundances were (**Fig 3A**).

**Figure 3:**
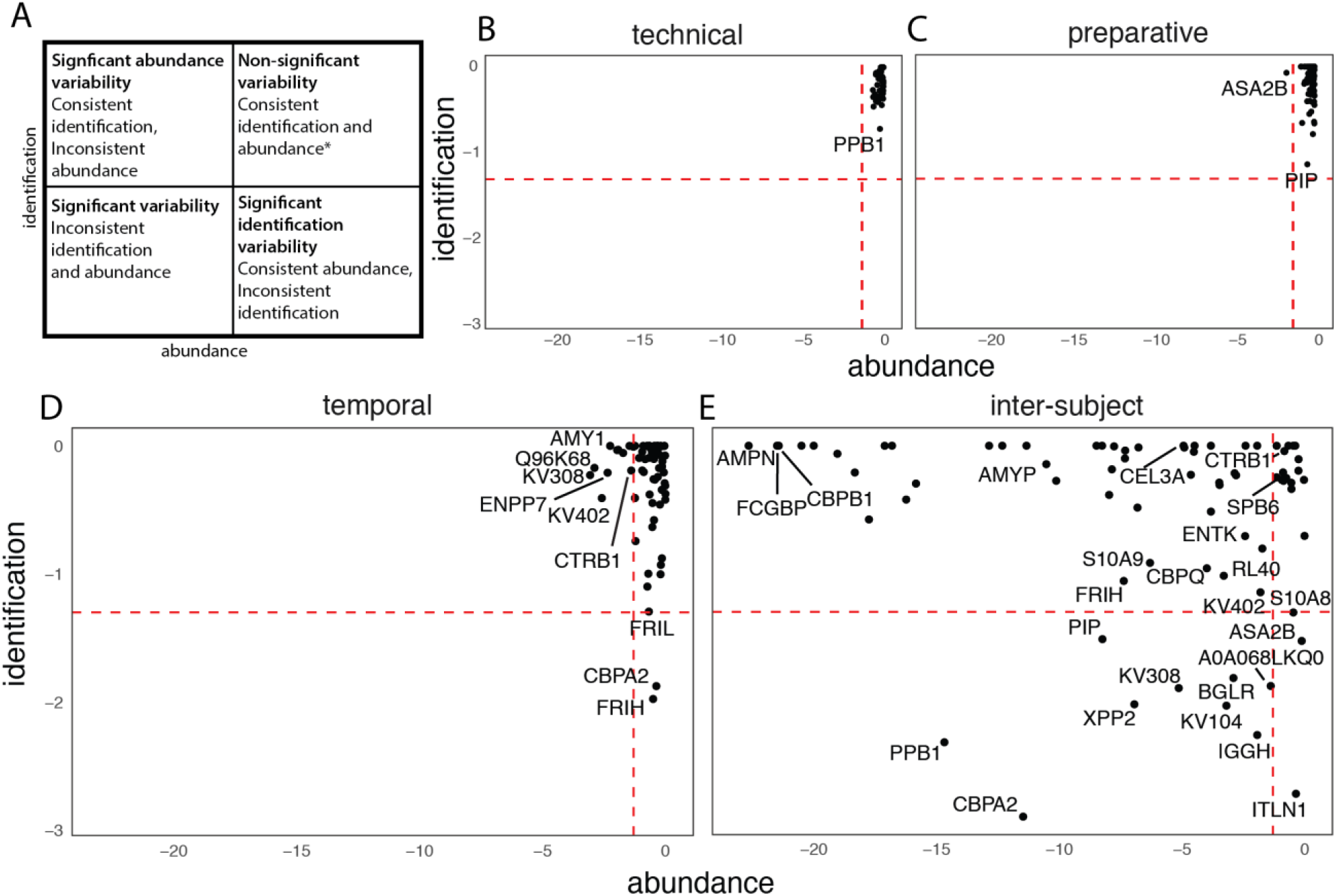
Variability significance testing using binary and conditional positive tests. **A**. Infographic to describe the corresponding significance with each quadrant. The x axis represents abundance variance (log(p)), determined by an ANOVA test. The y axis represents identification variance (log(p))– whether a given protein is identified or not – the significance of which was determined by a logisitic regression. The bottom left quadrant represents proteins that are significantly variable in both tests: these are proteins that are neither consistently identified nor consistently abundant among positive identifications. The top right quadrant represents a protein that is not significant: these are proteins that are consistently identified and measured with similar abundances. The other two quadrants represent proteins that are significant in one test, but not the other. Significance cutoff was p < 0.05, or log(p) < −1.30. **Supplementary Table 5** provides specific examples of these categories. *Non-significance could reflect proteins which were consistently present or consistently absent or had a consistently large degree of variability such that no single observation would appear to be an outlier. **B**. There were no significant proteins in technical replicates. **C**. ASA2B (Putative Inactive neutral ceramidase B) was inconsistently abundant in preparative replicates. **D**. 13 proteins demonstrated significant temporal varation. **11** proteins were inconsistently abundant and 2 proteins were inconsistently identified. **E**. 58 proteins were significantly variable for inter-individual variation. 9 proteins were inconsistently identified and abundant.

Non-parametric significance testing can be achieved with a logistic regression to determine whether a given protein was consistently identified (i.e., presence/absence). ANOVA provides comparable statistical rigor for evaluating how consistent protein abundance measurements are. However, both tests require that the number of proteins to be considered are less than the overall number of observations (i.e., N<160). We therefore selected a small subset of frequently identified proteins which were shared among all five subjects, and which would meet the ANOVA criteria. To meet these constraints we partitioned our protein identifications in three ways (**Fig. 2C**): those identified at least once (shared) in all subjects; the top 20% most abundant proteins (summed across all specimen in the dataset), and the top 20% with the greatest summed Shannon entropy, i.e., those proteins with the most similar abundances across the entire dataset. We found 74 proteins (estimated 0% FDR) that satisfied these criteria and were more enriched proteins relevant to the gastrointestinal tract, as compared to the entire dataset (**Fig 2B**). It is reasonable to expect that these shared proteins should be found in other healthy human subjects in future studies.

We subsequently tested each of these 74 proteins for four different types of variability: technical, preparative, temporal, and inter-subject as defined below. Each protein was assigned a p value from both a logistic regression (presence/absence) and ANOVA (abundance) tests, resulting in four state combinations describing whether a protein was inconsistently identified (presence/absence was significantly variable) or inconsistently quantified (significantly variable given that it was identified) (**Fig 3A**).

### 2.3 Instrument and operator error are largely insignificant sources of fecal proteome measurement variance

The stochastic nature of “shotgun” tandem mass spectrometry experiments (16) could limit our ability to identify and quantify stool proteins across multiple specimens. To evaluate this source of variance with respect to other experimental or biological sources, we compared identification rates and protein quantifications from independent replicate stool preparations (preparative replicates), and from sequential LC-MS analyses of each preparation (technical replicates) (**Fig. 1B**). None of the 74 proteins demonstrated significant variability among technical replicates with respect to their ability to be identified (general linearized model); p <0.05, log(p) <−1.3), or quantified (when identified) (ANOVA; p <0.05, log(p) <−1.3) across all subjects and time points (**Fig 3B**).

We expected preparative variation to be greater than technical variation due to the combined effects of intra-specimen heterogeneity, human error, and drifting LC-MS conditions over time. One protein, ASA2B, an intracellular ceramidase highly expressed in the gastrointestinal tract, demonstrated abundances which varied significantly between preparative replicates (abundance log(p) = −1.70), despite being consistently identified (identification log(p) = −0.07) (**Supp Fig 4**). This result could suggest that ASA2B may have been disproportionately distributed within each stool specimen – lending to its inconsistent identification. However, datasets considering thousands of measurements often have statistical outliers, which could be another explanation for our quantification variance of ASA2B. Overall, we conclude that preparative variation was not a substantial factor in these proteins’ measurement.

### 2.4 Temporal variances among stool proteins were significant and had no clear directional trends

Prior analyses of human stool proteins gave limited consideration to longitudinal variation in the absence of a perturbation (8, 10, 17). Accordingly, the extent to which stool proteins vary in healthy adults over time within subjects is unclear. Our experimental design (**Fig. 1A**) allowed us to examine the temporal variability of host stool protein abundances and identifications.

Seventeen percent (13/74) of the shared protein subset demonstrated significant variation in identification (2) or abundance (11) over time (**Fig. 3D**), which contrasts with just one inconsistently identified protein across all preparative replicates. We note that over half of the proteins found to significantly vary in abundance over time had immunoglobulin-related annotations. The shared protein subset is already significantly enriched in immunoglobulin-related proteins, and this finding supports that specific immunoglobulin protein levels can fluctuate over time in the GI tract, as was recently reported by Tropini et al (18).

In contrast, we expect proteins that are induced by inflammation to be consistently expressed over time within healthy subjects. Accordingly, we found that the identification and abundances measured for the neutrophil-expressed inflammation marker, calprotectin (S100A9) did not vary significantly over time (log(p) > −1.30 by both identification and abundance tests). Two other known markers of gastrointestinal inflammation within our shared protein subset were consistently identified and quantified over time in our assay: DMBT1, which was shown to be upregulated in Crohn’s disease (19), along with S100A8, another fecal calprotectin subunit (**Fig. 3D**). However, there were many more clinical biomarkers in our gastrointestinal proteome that were excluded due to their lack of shared presence or high abundance or entropy amongst subjects. Current clinical biomarkers include S100A12, lactoferrin, metalloproteinases, myeloperoxidases, and neutrophil elastase, all of which we observed in our full dataset. The temporal variances we observed across the shared protein subset (**Fig. 3D**) could reflect natural variation in a donors’ diet and lifestyle. To evaluate this, we examined all proteins in the data set (**Fig. 4A**). If the stool proteins were to match previous microbiome results, we would expect to see temporal variability unique to each individual in unperturbed, uncontrolled diet conditions (14, 20). The resulting principle component analysis (PCA) with Bray Curtis distance metric indicated that each subjects’ stool proteins varied temporally to different extents, confirming that stool protein temporal variability mirrors previously described temporal microbiome variability. To further quantify this aggregate variance, we measured the Euclidean distances between each time point in this PCA space. We found that sequential days were no more similar than days separated by a week or a month. Furthermore, we found that stool protein profiles did not demonstrate any obvious progression over time in any subject (**Fig 4C**). Furthermore, we found that the summed Euclidean distances between each point also ranged substantially across each individual. This temporal sampling helps to establish a range above which a researcher could describe as a perturbation.

**Figure 4:**
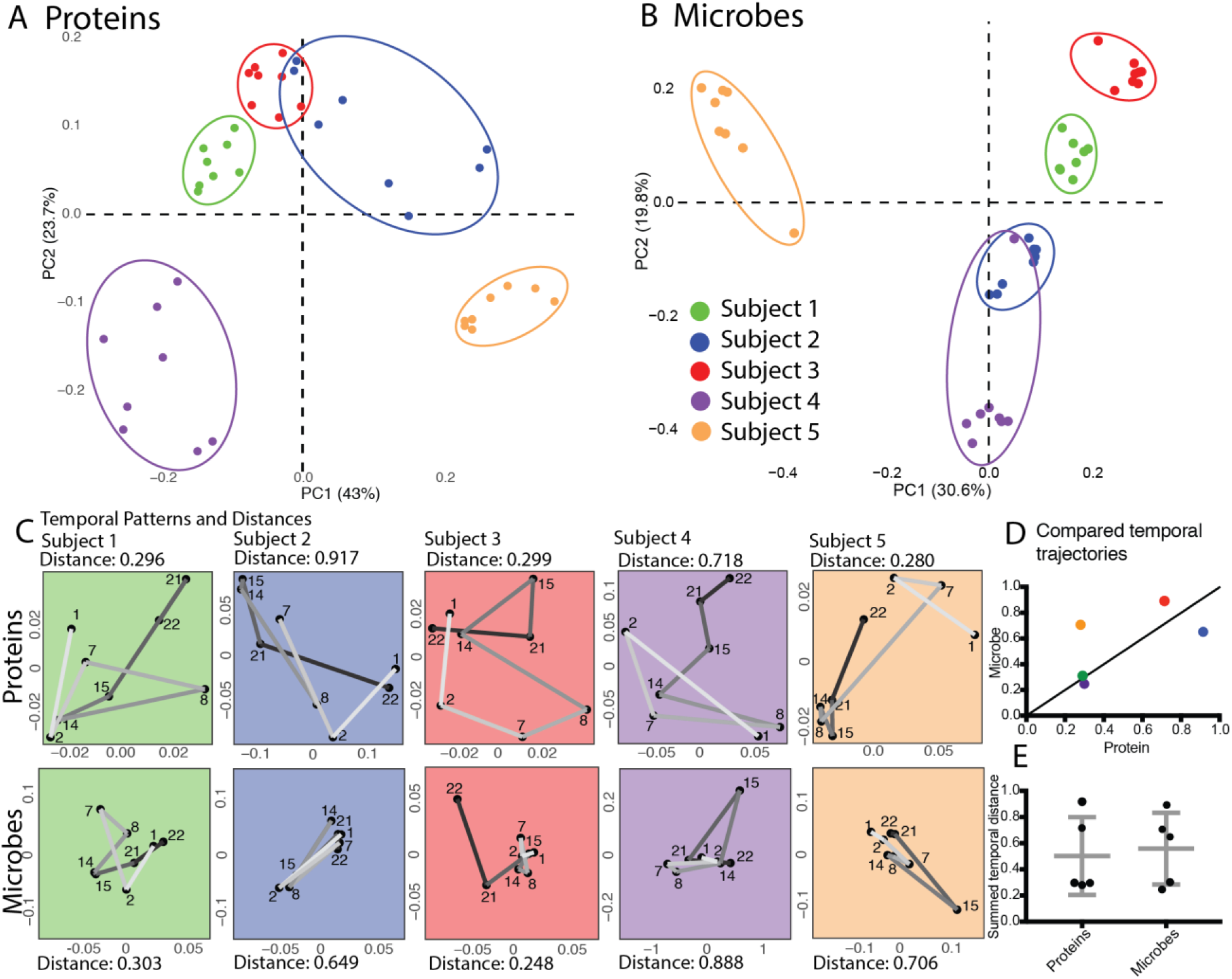
Stool proteome and microbiome show inter-individual variance and no temporal trend. A, B Principal coordinate analysis using Bray-Curtis dissimilarity for both proteins and microbes to examine inter-individual and temporal trends See Sup Fig 6 for ordination with additional distance metrics. **A**. Data from all proteins (456). Subject clusters are evident, but overlapping. Subjects 4 and 5 are most separate along the 1^st^ PCoA axis; subject 3 is most separated from both 4 and 5 along the 2^nd^ axis. **B**. Data from 1213 microbes meeting or surpassing both 5% presence and 10 count abundance. Subject clusters are almost entirely distinct with one exception noted below. Subjects 3 and 5 are most separated along the 1^st^ axis, and subject 4 from both 3 and 5 along the 2^nd^ axis. A single specimen (subject 4, day 15) clearly clusters as expected with the subject’s other time points along the 3^rd^ principal coordinate axis (15% of variability), which is not shown in Fig 4B. **C**. Temporal distances and patterns in the protein and microbe data sets, based on Bray Curtis PCoA vectors from (A) and (B) that have been zero-centered for each subject separately. Numbers indicate sequential specimens; temporal pattern is represented as a gradient from white to black. No temporal trends are apparent in either data set. Temporal distance is the sum of sequential point-to-point (Euclidean) distances. Sum allows us to quantify “movement.” Larger sums equate more movement and variation in time. **D**. Protein and microbial temporal distances are not correlated across subjects (r^2^ = 0.35). Black line indicates a 1:1 microbe to protein temporal distance ratio. **E**. Temporal distance is similar for proteins and microbes.

Noting that eight proteins accounted for almost 50% of total protein abundance across all specimens and replicates, we wondered if any could have had a disproportionate influence on our temporal measurements. Accordingly, we compared how these protein (CTRC, CEL3A, AMY1, CEL2A, CEL3B, A1AT, TTHY, IGKC) abundances change over time within each donor and found that none accounted for the global temporal distances we observed across every subject (**Fig 4C, Supp. 5**). For example, we found that subjects two and four traversed the largest distances in PCA space over time (0.917 and 0.718, respectively **Fig 4C**). However, proteins such as A1AT (20% and 3.3% of total abundance on average, respectively) fluctuated widely across most time points measured from subject 2 (e.g., a max 3.4 fold change between any two days) but were fairly consistent in subject 4 (at most a 1.6-fold change between any two days) (**Supp Fig 5**). We conclude that the collective, and largely distinct profile of each subject’s stool proteomes accounted for dataset-wide temporal instability, rather than the most abundant proteins.

Proteins that greatly fluctuate over time during states of health have diminished diagnostic utility. Although we note substantial temporal variance for most proteins we measured from these five subjects, our assay also identified proteins that were largely stable over time. For example, one trypsin (TRY1, the 12^th^ most abundant protein when summed across all specimens) showed no significant identification and abundance variability over time (abundance log(p) = −0.75 and identification log(p) = 0.00). This protease was previously shown to increase substantially with inflammation stressors (21). Knowledge of stable proteins like these proteases can help to distinguish subjects that are experiencing gastrointestinal inflammation.

### 2.5 Stool proteins vary significantly between healthy human subjects

We previously showed that host-expressed stool proteins could distinguish healthy humans from each other (n=3) (8). However a subsequent study of 29 subjects showed only slight ability to distinguish obese from non obese individuals based on 41 proteins that were measured and quantified by different methods than those described here (10). Both of these experiments only studied specimens collected at a single time point for each subject. With the aim of distinguishing subjects using multiple temporal measurements of host-expressed [rpteins, we considered how individual proteins varied between subjects.

Almost 80% (58/74) of the shared protein subset demonstrated significant variation in identification (3), abundance (46), or both (9) between all subjects (**Fig. 3E**). There were 48 more proteins varied significantly between subjects (**Fig 3E**) than within individual subjects over time (**Fig 3D**). This finding allowed us to conclude that the largest source of variance in host protein expression was inter-subject versus temporal, preparative, or technical variance. Further, inter-subject protein abundance varied 7-fold more than temporal abundance (minimum temporal log(p) = −3.07 for protein KV308; minimum inter-subject log(p) = −22.65 for protein AMPN). Calprotectin subunit S100A9’s abundance variability was exemplary of this larger inter-subject variation trend (inter-subject log(p) = −6.30, temporal log(p) = −0.92) (**Supp. Fig. 4**). While fecal calprotectin (heterodimer of S100A9 and S100A8) levels are used to distinguish between healthy and inflamed guts, our experiment suggests that S100A9, despite little temporal variability within subject, has a large intrinsic abundance variability between healthy subjects (quantification log(p) = −6.30 identification log(p) > −1.30) and S100A8 was consistent by quantification and identification (both log(p) > −1.30) when assessed through this LC-MS/MS platform.

Overall, there were 48 proteins with only inter-subject variability (significant identification or abundance variability) and 3 proteins with only temporal variability (**Supplemental Table 1**). The largest source of variability in this shared protein subset is protein abundance among healthy people.

While we have shown that proteins varied significantly across all subjects, it does not necessarily follow that these proteins could distinguish subjects from each. We therefore evaluated the extent to which host stool proteins were distinct per individual by considering all of the proteins we reported in Figure 2 and not just the shared protein subset (See Fig legend 4A). We observed strong, though not complete, inter-subject grouping (Fig 4A). We used Bray-Curtis dissimilarity to be consistent across both proteome and microbiome data types, but Euclidean distance is often applied to protein data sets as a measure of specimen dissimilarity (which, unlike Bray-Curtis, considers the shared absence of a protein in specimens as evidence of their similarity) (**Supp Fig 6**). The Euclidean distance separated subjects with a similar distinction to the Bray Curtis separation.

Subject 4 was the most distinct individual, using the Bray Curtis distance (**Fig. 4A**). Several proteins were upregulated in this individual relative to the others, including CUB/zona pellucida-like domain-containing protein (CUZD1), pancreatic zymogen molecule 2 (GP2) (**Supp. Fig. 7**). Both of these proteins are targets of pancreatic autoantibodies and GP2 is upregulated in Crohn’s disease patients (22).

It would be useful to identify a set of proteins, which are commonly found in healthy people, with a defined, narrow abundance range. In our shared protein subset, we found 16 proteins that were consistently identified and quantified across all people. This set of proteins has a range of attributes, functions, and identities including disulfide bonds, serine protease inhibitors and pancreatic digestive enzymes like chymotrypsinogen B1 (CTRB1) (Fig 4E). These could be potential candidates for tracking “return to health”, as they are observed and quantified consistently over time and between people.

However, most proteins we measured did vary between subjects. This could reflect differences in the gut microbiota, which is known to have individualized compositions and interact with the host, affecting host protein secretions (8, 23, 24).

### 2.6 Studying proteins and microbes together can elucidate orthogonal conclusions

As with host proteins, and consistent with expectations (23, 25, 26), the taxonomic composition of the gut microbiota was more similar within a subject over time than it was between subjects (p<0.0001) (**Fig 4B, Supp Fig 8**). Subjects appeared equally distinct from one another, with respect to their microbiota and their stool proteins when comparing PCAs (**Fig 4A,4B**). This might be surprising given that the human gut microbiota is comprised of many independent species, while stool proteins are derived from the genome of a single host. However, one might also expect pronounced patterns of inter-subject co-variation between microbe and protein data sets: host-expressed proteins and the microbiota may directly interact within a host’s gut. Furthermore, both stool proteins and gut microbiota are influenced by the same intrinsic (e.g., inflammation status) or extrinsic (e.g., diet) factors, which tend to vary between hosts. Accordingly, we note that the same three subjects (3, 4, and 5) were furthest from one another according to the ordinations of **Fig 4A,B** for both proteins and the microbiota. However, such comparisons were inconsistent within this small set of subjects: subjects 2 and 4 demonstrated similar gut microbiota composition (with the exception of one outlier day in subject 4) (**Fig. 4B**), but showed high variance among host stool proteins (**Fig. 4A**). Additionally, other subjects were inconsistently distinguished by proteome and microbiome data sets: proteins from subject 2 partially overlapped with those from subject 3, but the gut microbiota completely separated these individuals.

Using PCA coordinates, we extracted both temporal patterns and Euclidean distances between days and subsequently summed these temporal distances (**Fig 4C**) alongside temporal microbial family changes (**Supp Fig 3**) and, as we observed with the proteins, no single family or even phylum was responsible for explaining overall temporal patterns. For example, subject 4, whom had the greatest temporal distance (0.888, **Fig 4C**), was characterized by a large shift in *Bacteroidetes*, especially on day 15. However, subject 5, whom had the second largest temporal distance (0.706, **Fig 4C**), was characterized by large shifts of *Verrucomicrobiaceae*, a microbe family in a completely separate phylum. We conclude that temporal distances for individuals were driven by subject-specific temporal microbial shifts.

Temporal patterns were also quite dissimilar between protein and microbial data sets (**Fig 4C**). For example, the day 15 and 22 specimens from subjects 3 and 4 were quite distinct from other days, as assessed from the microbiota. However, these particular days were quite typical of the subject as assessed from their stool proteins. We found that proteins and microbes exhibited little coordination over the duration of a month. Furthermore, we note that the ordinations plotted here depended greatly on the choice of distance/dissimilarity metric. We used Bray-Curtis dissimilarity to be consistent across both data types, however microbiome researchers often compare specimens using methods that account for the phylogenetic relatedness of microbial taxa (e.g., weighted Unifrac), unlike Bray-Curtis. Ordinations using these metrics were rather different from Bray-Curtis (**Supp Fig 6**). However, they did not suggest any consistent inter-subject variability patterns shared between stool proteins and the gut microbiota (**Supp Fig 6**). We conclude that microbe and proteome-focused assays capture related, yet distinct aspects of the dynamic gut ecosystem.

### 2.7 Stool proteins detect inflammation signatures

One distinct aspect of the protein profile is the ability to reveal known host inflammatory features, due to the annotation-rich and disease-focused nature of human protein databases. For example, both calprotectin subunits distinguished subject 2 from the other four subjects (**Supp Fig 7**). We therefore considered whether any subjects in our study could be distinguished by the presence of other known inflammation-associated proteins, and whether proteins with other functional annotations mirrored these inflammation-related patterns. We selected proteins enriched in subject 2, but sparse in all other subjects and found that they were enriched in proteins known to be associated with inflammation like MHC, immunoglobulins, neutrophil elastase, and serine protease inhibitors (**Supp Fig 9**).

To this answer this issue further, we selected quantification data for known fecal clinical biomarkers from the entire dataset. This set included lactotransferrin, calprotectin S100A9 and S100A8, neutrophil elastase, and myeloperoxidase (27). With the exception of S100A9, these proteins were significantly more abundant in subject 2 compared to most subjects (**Fig 5**), consistent with an elevated basal inflammation level. Myeloperoxidase and neutrophil elastase were significantly enriched in subject 2 compared to all other subjects (p<0.05 and p <0.05). It is possible that subject two is slightly inflamed- either clinically or, more likely, within an overall normal range in the healthy spectrum.

**Figure 5:**
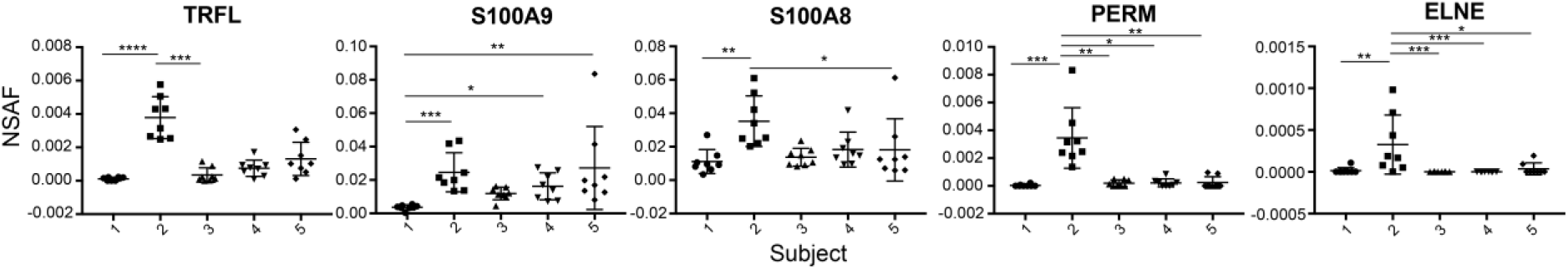
Known clinical biomarker protein abundance in the stool proteome suggests an inflammation signature in Subject 2. These proteins are known clinical biomarkers (27). X axis shows corresponding subject, y axis shows normalized abundance (NSAF). Statistics were run using a kruskill-wallis anova with Dunn’s multiple comparison test. P values from Kruskal-Wallis ANOVA test are denoted as such: * <.05 **<.005 ***<.0005 ****<.00005. Proteins are labeled as graph title. Subject 2 is significantly upregulated compared to all other subjects in PERM and ELNE.

## 3. Discussion

We report here a baselining investigation into human stool protein variation with respect to technical and preparative replicates, multiple samplings over time, and between people. We observed that mass spectrometry-based identification and quantification of stool proteins provided consistent values for technical and preparative replicates. Of note, individual stool protein measurements were observed and quantified with significant variance over time and were most variable between people. Stool proteins distinguished healthy subjects from one another, in a manner complementary to 16S rRNA gene sequences. We believe this study helps to establish a useful foundation for examining how stool proteins may provide insight into a wide range of enteric diseases. We selected proteins with the highest entropy, that were identified amongst all subjects, and were most abundant, which biased against the identification axis in our probability plots: most of the 74 proteins should have been identified consistently. The features we learned about time and inter-subject variance when we expanded to the full protein set supported our results and suggests that, despite bias, our results are reflective of the human gastrointestinal proteome as a whole. While confidence values can speak to the overall variance of a protein, our variability readouts using nested linear models allowed us to determine the nature and significance of variability for each protein.

We suggest that human stool proteins can provide an important complement to microbial taxa-based assays to better diagnose and classify human gastrointestinal health. Data from both stool proteins and the microbiota will be more informative than either alone. Some microbial strains have shown to be inflammatory in one subject and benign in another subject (28). Perhaps by analyzing the microbes and proteins in complement, the health insights obtained from both the stool proteome and microbiome can provide a fuller picture for health status.

More work is needed to scope out the full range of health. Larger studies of healthy humans will allow more rigorous techniques like cross validation between patient cohorts, and a fuller picture of the spectrum of healthy human stool protein profiles. In the future, stool proteins could be used to track inflammation, but also a patient’s return to health. Longer longitudinal studies over months could inform researchers of the natural temporal dynamics of these analytes. Additionally, studying healthy subjects versus subjects with various forms of intestinal disease will allow researchers to curate stool proteins potentially diagnostic of gastrointestinal health.

## Supplemental Figures

**1:**
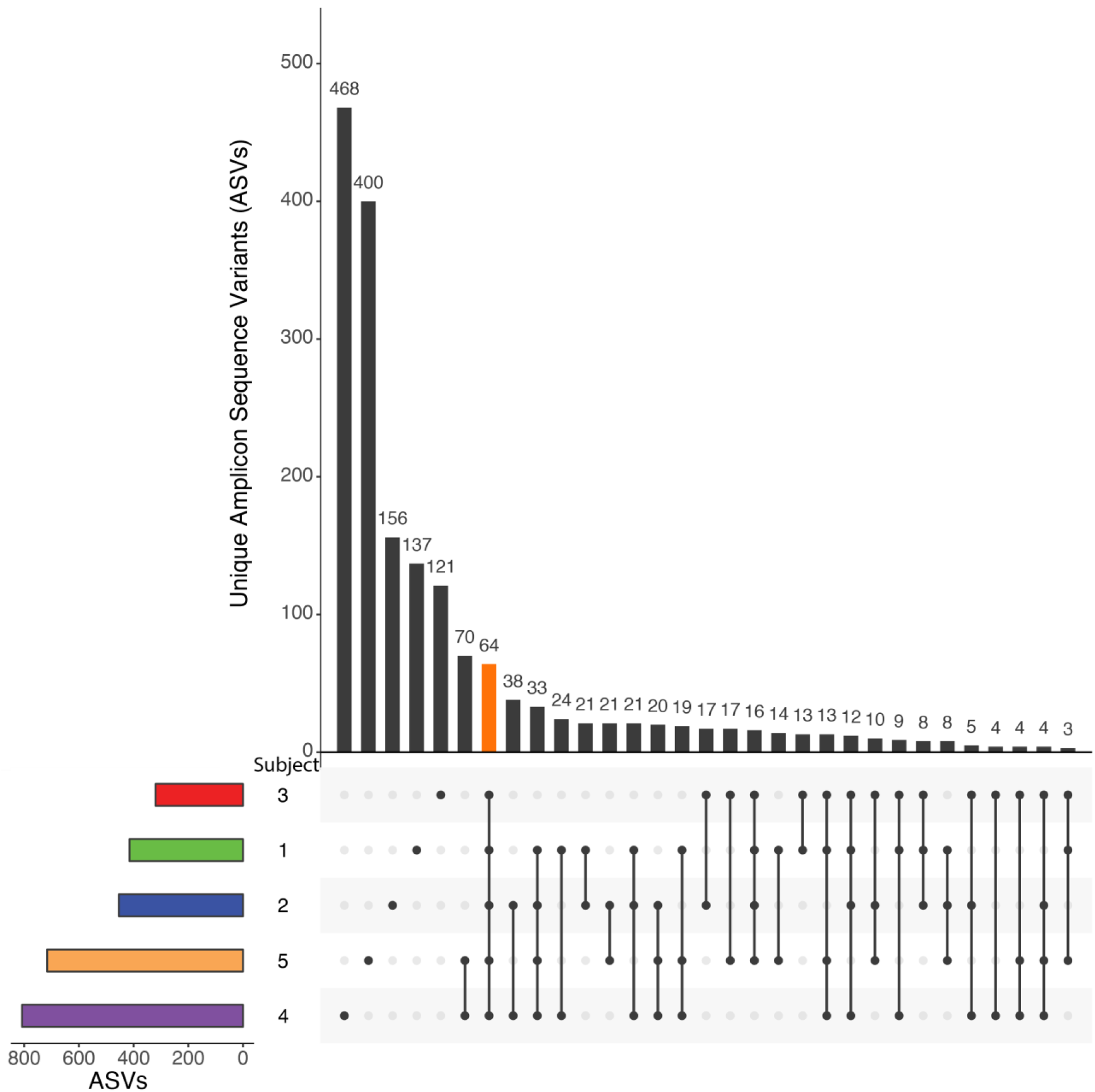
Majority of microbes are identified in only one subject than shared across multiple subjects. Subset plot showing the detection of ASVs (Amplicon Sequence Variants) across subjects. Of 1,770 ASVs detected, 1,282 ASVs (72%) were identified in only one subject; 64 ASVs (4%) were found in all subjects..

**2:**
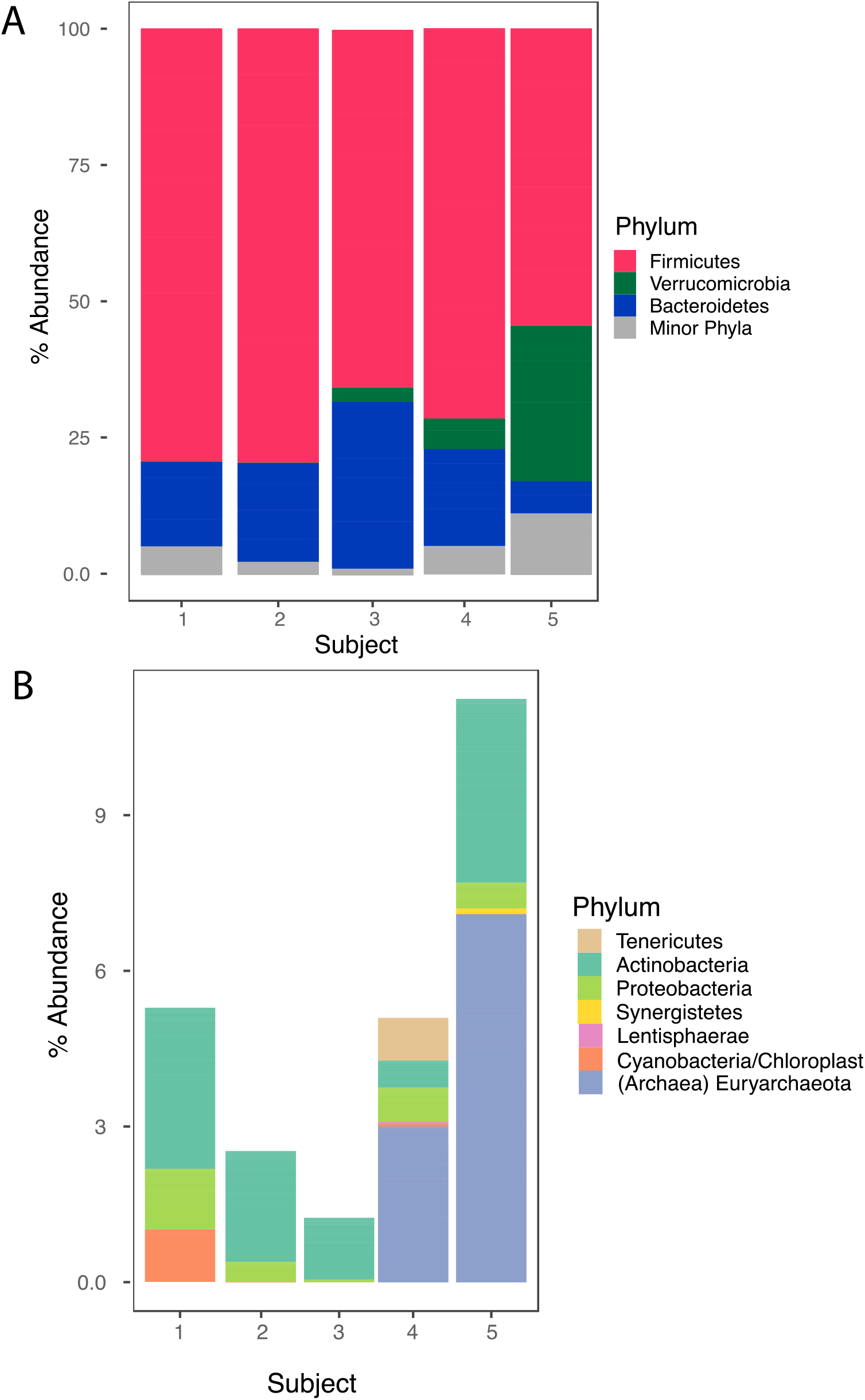
Firmicutes and Bacteroidetes phyla dominant in all subjects. Stacked bar plots display average percent abundance of phyla per subject for high abundance phyla in A and low abundance phyla in B. The abundance of Verrucomicrobia in subject 5 is atypical, but patterns of variability between subjects and over time (Sup.Fig 5) are consistent with biological variability rather than contamination or PCR bias.

**3:**
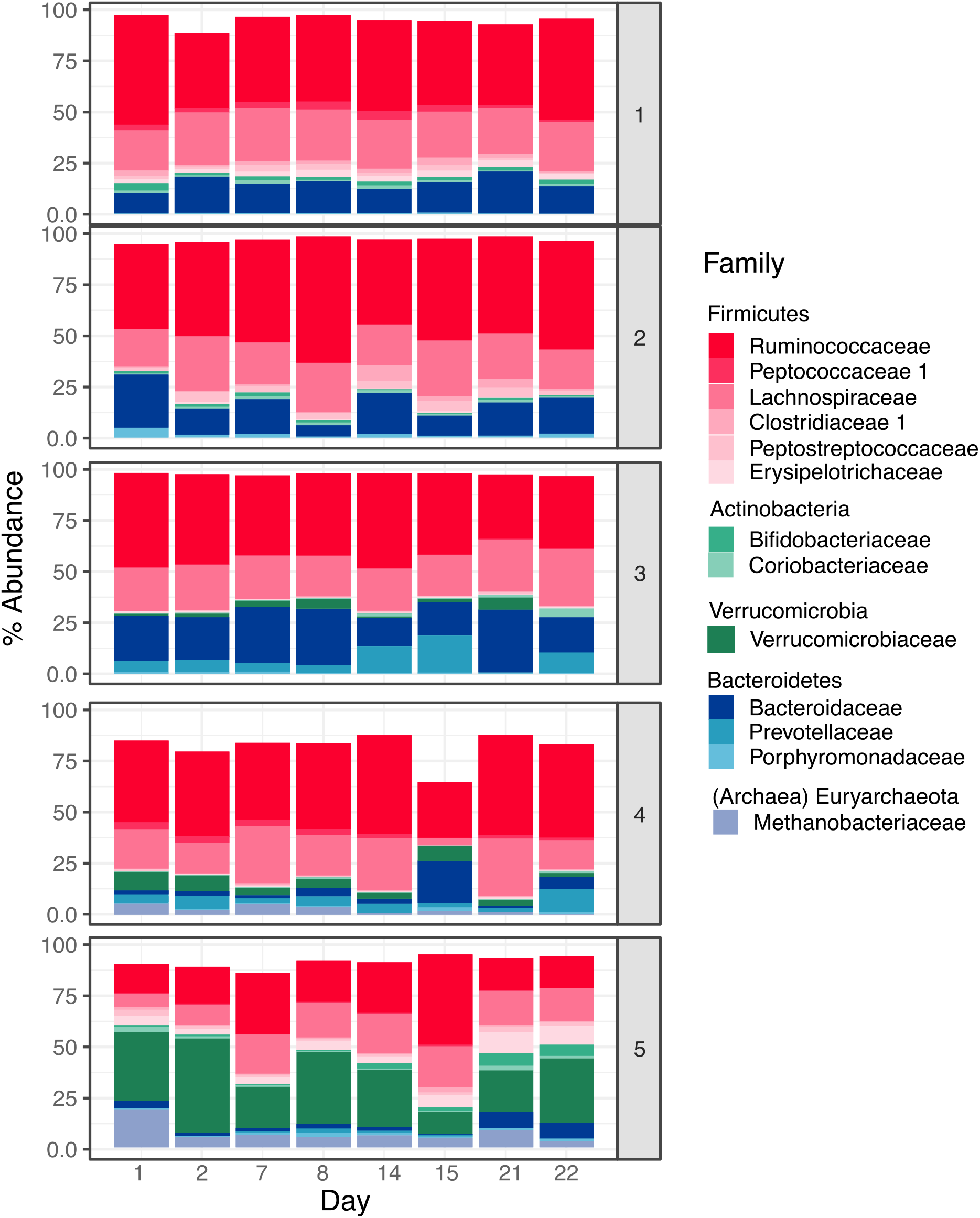
Temporal variability of microbial families. Stacked bar graphs show the percent abundance of 13 families in each subject over time. The distinct set of families present in each subject persists, with varying abundances over time.

**4:**
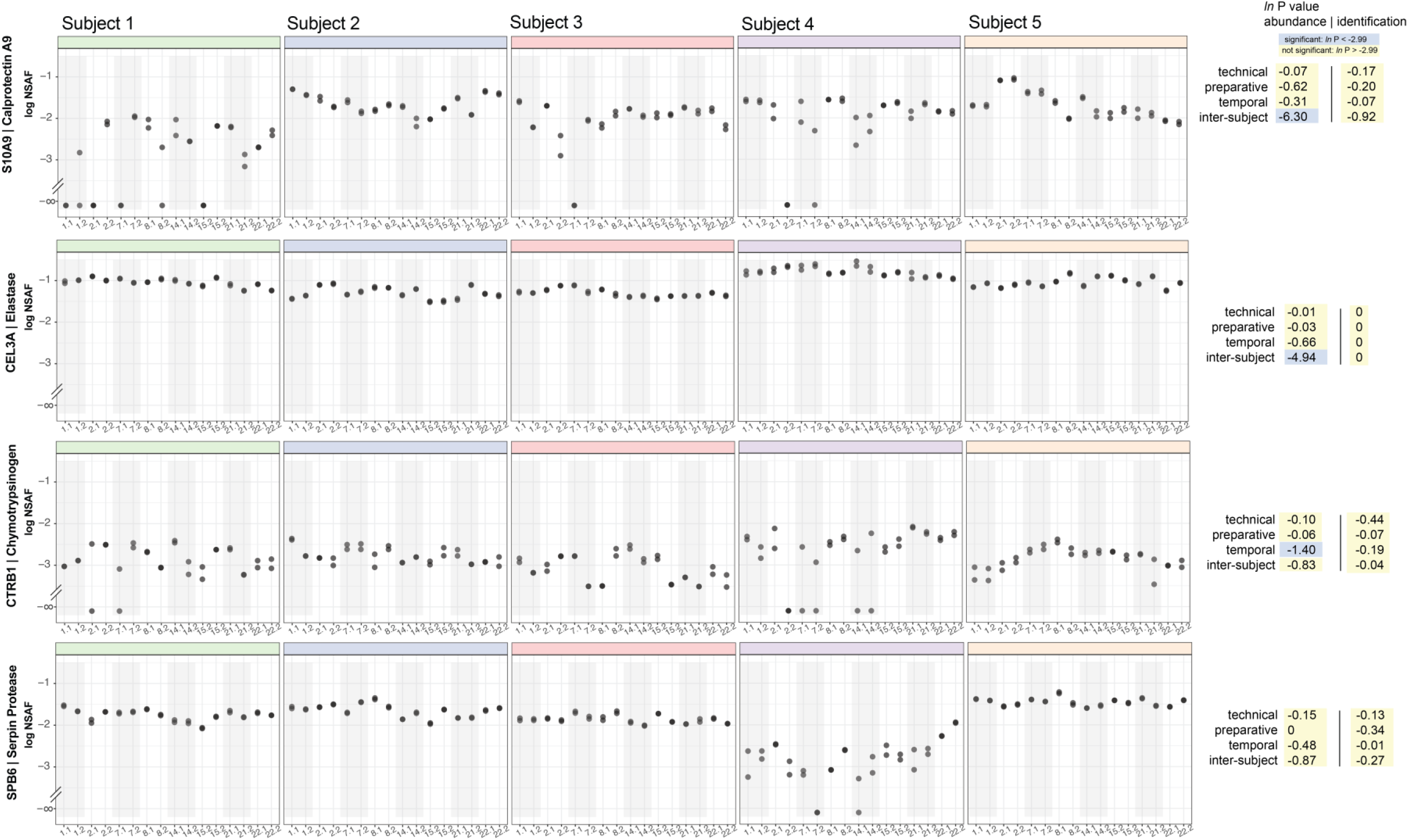
Variability is not contingent on only abundance. Specimens are labeled with regard to day (1–22) and preparative replicate (either nothing, or .2). Each plot is a different subject’s respective protein abundance over time and preparative replicate. log (NSAF) abundance values are plotted on the y-axis. Calprotectin (A) showed significant inter-subject variance. We additionally picked three proteins that represented a range of abundances: a high abundance protein (CEL3A, B), a low abundance protein (CTRB1, C), and a protein with matched abundance to calprotectin (SPB6, D). Calprotectin and CEL3A both had only inter-subject variability, and CTRB1 had only temporal variability.

**5:**
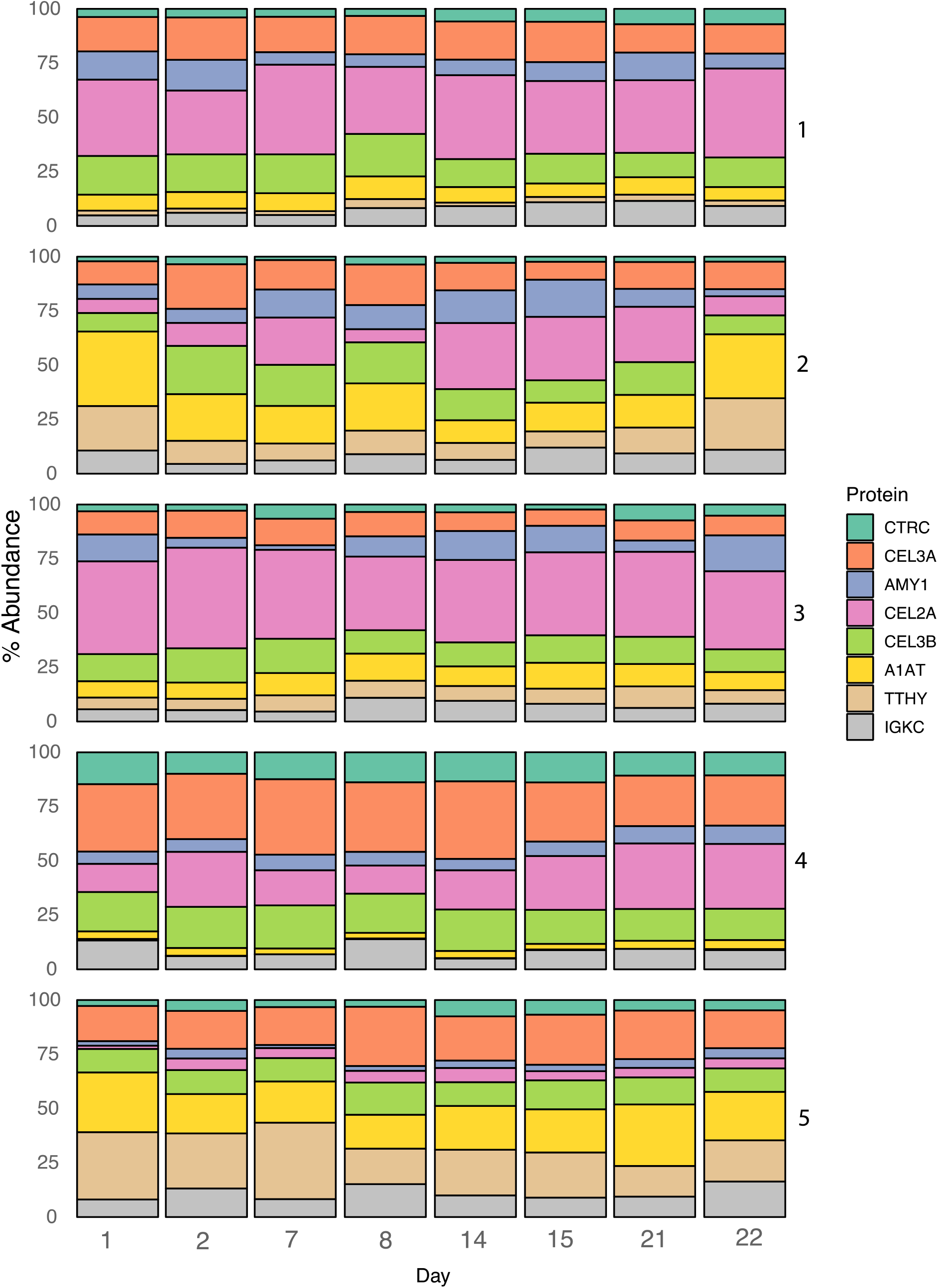
Top 8 most abundant proteins contrast in abundance and temporal variation between subjects. Stack bar graphs show % abundance protein changes per subject over time. There are no obvious protein abundance change trends over time across all subjects or in a single subject. This supports the conclusion that there are no directional temporal trends in stool proteins over time.

**6:**
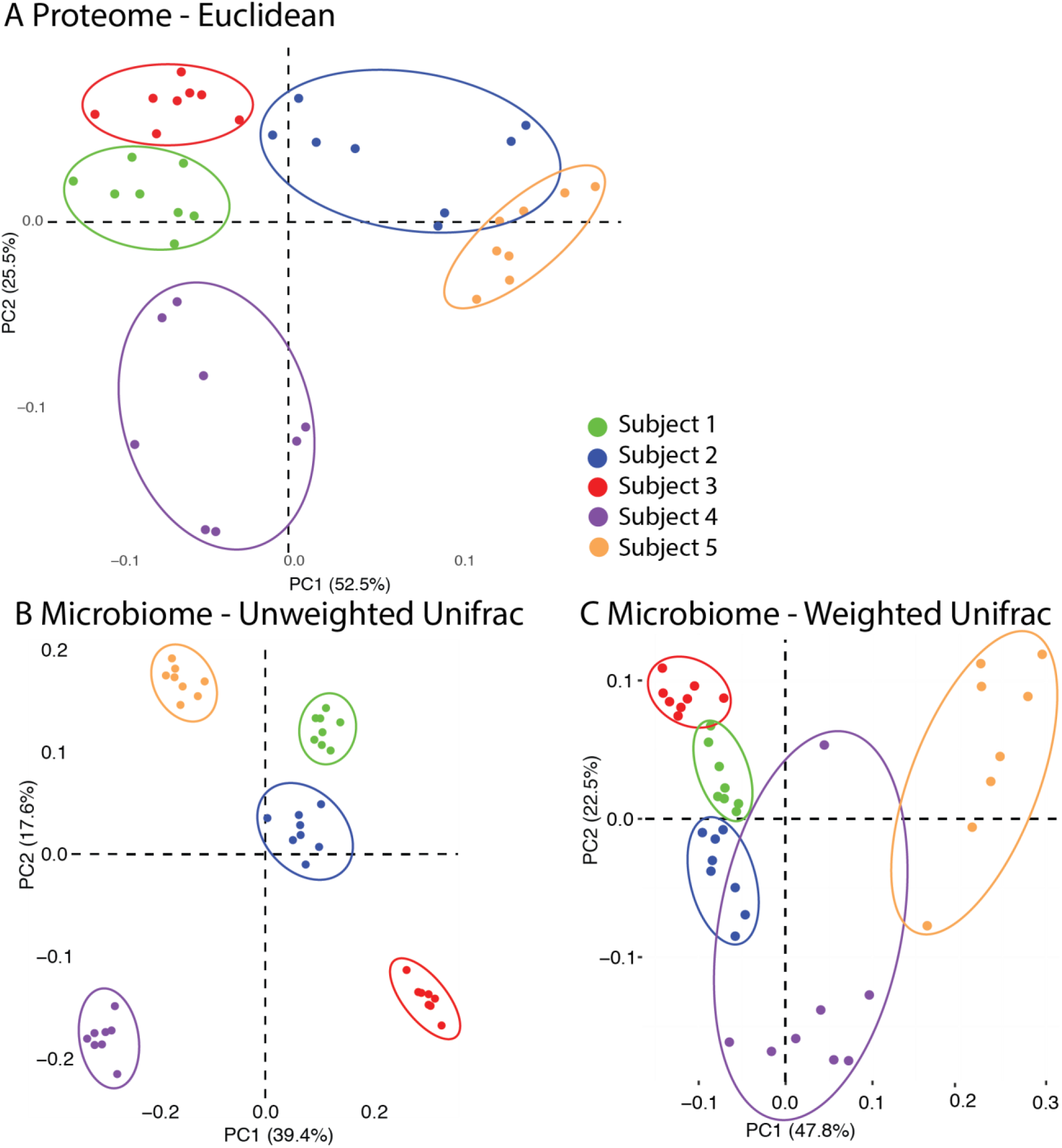
Different patterns appear with ordination using different dissimilarity metrics. Supplemental ordination plots corresponding to Fig 4A&B. Microbes and proteins were filtered in the same way, the only difference is the choice of distance metric. A) Protein ordination plot using Euclidean distance. Ordination differentiates subjects in a manner similar to Bray Curtis ordination. Subject may be slightly less differentiated, which could be due to the consideration of zero values to the dissimilarity. B) Microbe ordination plot using unweighted Unifrac distance, which accounts for the phylogenetic relatedness of microbial taxa but ignores abundance, considering only taxon presence/absence. Subject clusters are small and well-separated, reflecting different sets of taxa that persist in each subject. C) Microbe ordination plot using weighted Unifrac distance, which accounts for both phylogeny and relative abundance of microbes. Subject clusters are distinct, but less so than with unweighted Unifrac or Bray-Curtis (which considers abundance but not phylogeny), indicating that abundant taxa tend to be more closely related across subjects than is true for the microbiota as a whole. The dissimilarity of a single specimen from others within subjects 4 and 5 primarily involves changes in the relative abundance of taxa, not their identity.

**7:**
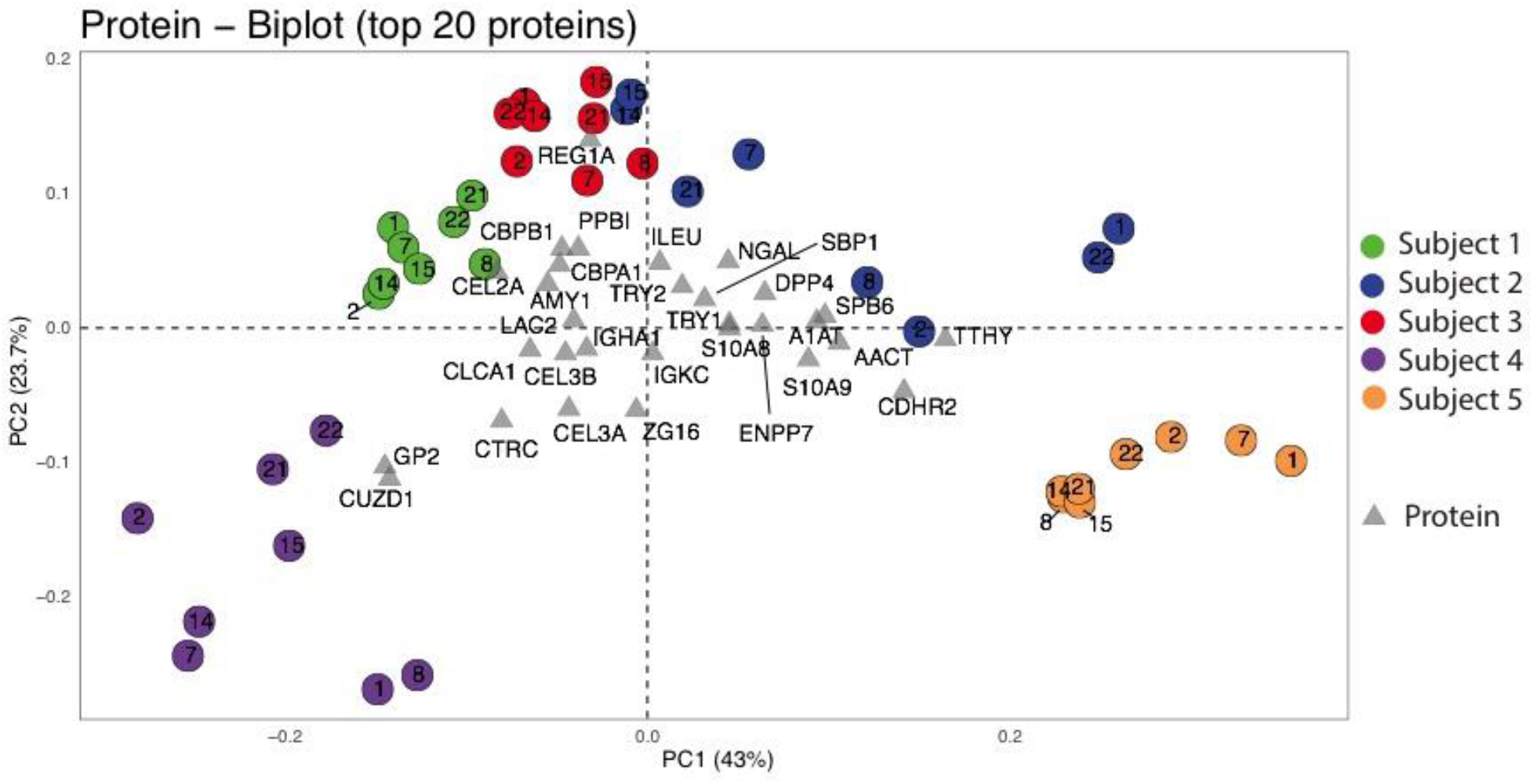
Proteins that separate subject 4 from other healthy humans include GP2 and CUZD1. Dimensions and directionality correspond to Fig 4A. Each circle represents a specimen: the color corresponds to the subject and the number corresponds to the day the specimen was acquired. Each protein is labeled, and location of label shows contribution directionality. Subject 4 was most distinct from the rest of the subjects due to comparative increased abundance of pancreatic zymogen molecule 2 (GP2) (GP2) and CUB/zona pullucila-like domain-containing protein (CUZD1). GP2 and CUZD1 are both targets of pancreatic autoantibodies.

**8:**
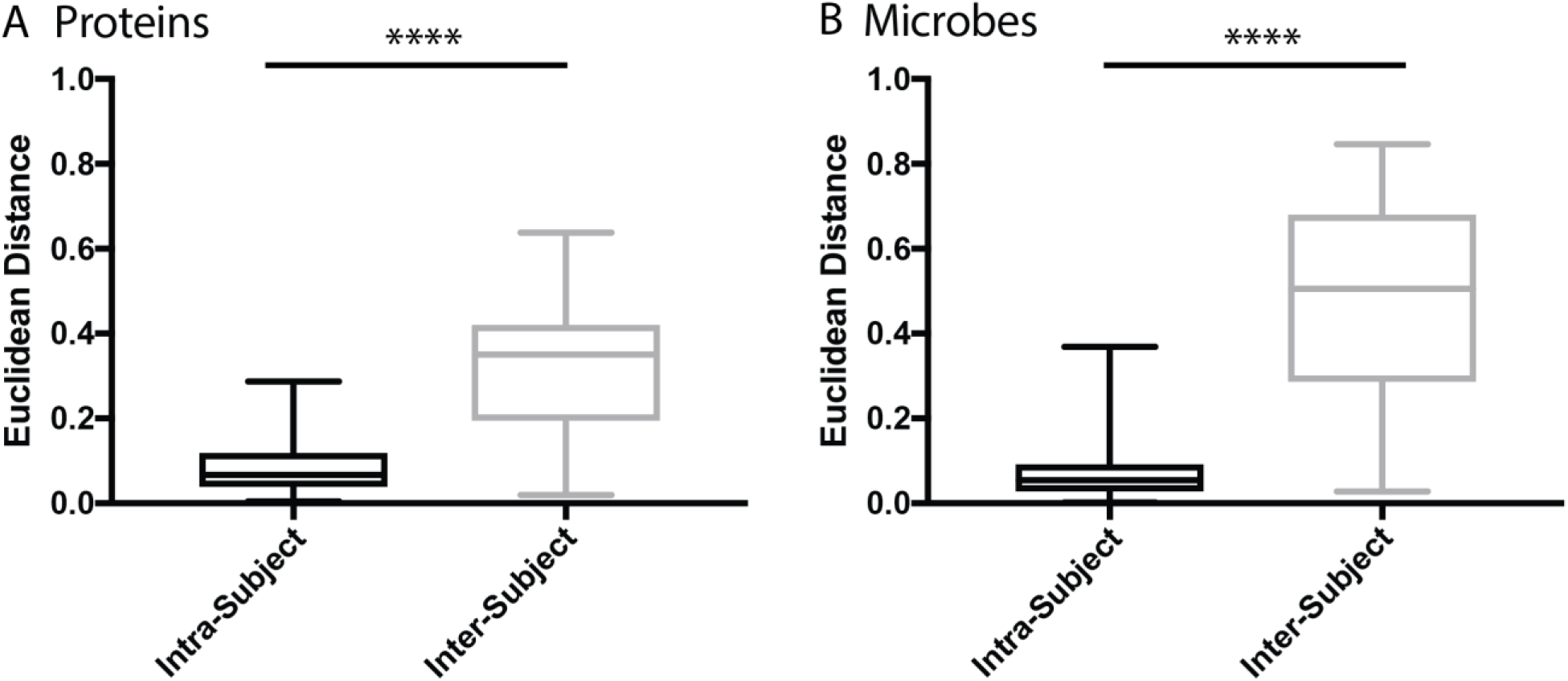
Protein and microbe Euclidean distances from PCA coordinates show intra-subject specimens have a smaller distance between each other than between inter-subject specimens. Euclidean distances between all coordinates were obtained from the PCA coordinates in **Fig 4A,B. A** Specimens from the protein PCA (**Fig 4A**) are more similar within a single subject (intra-subject) than between subjects (inter-subject) (p<0.0001, Mann-Whitney U). **B** Specimens from the microbe PCA (**Fig 4B**) are more similar within a single subject (intra-subject) than between subjects (inter-subject) (p<0.0001, Mann-Whitney U).

**9:**
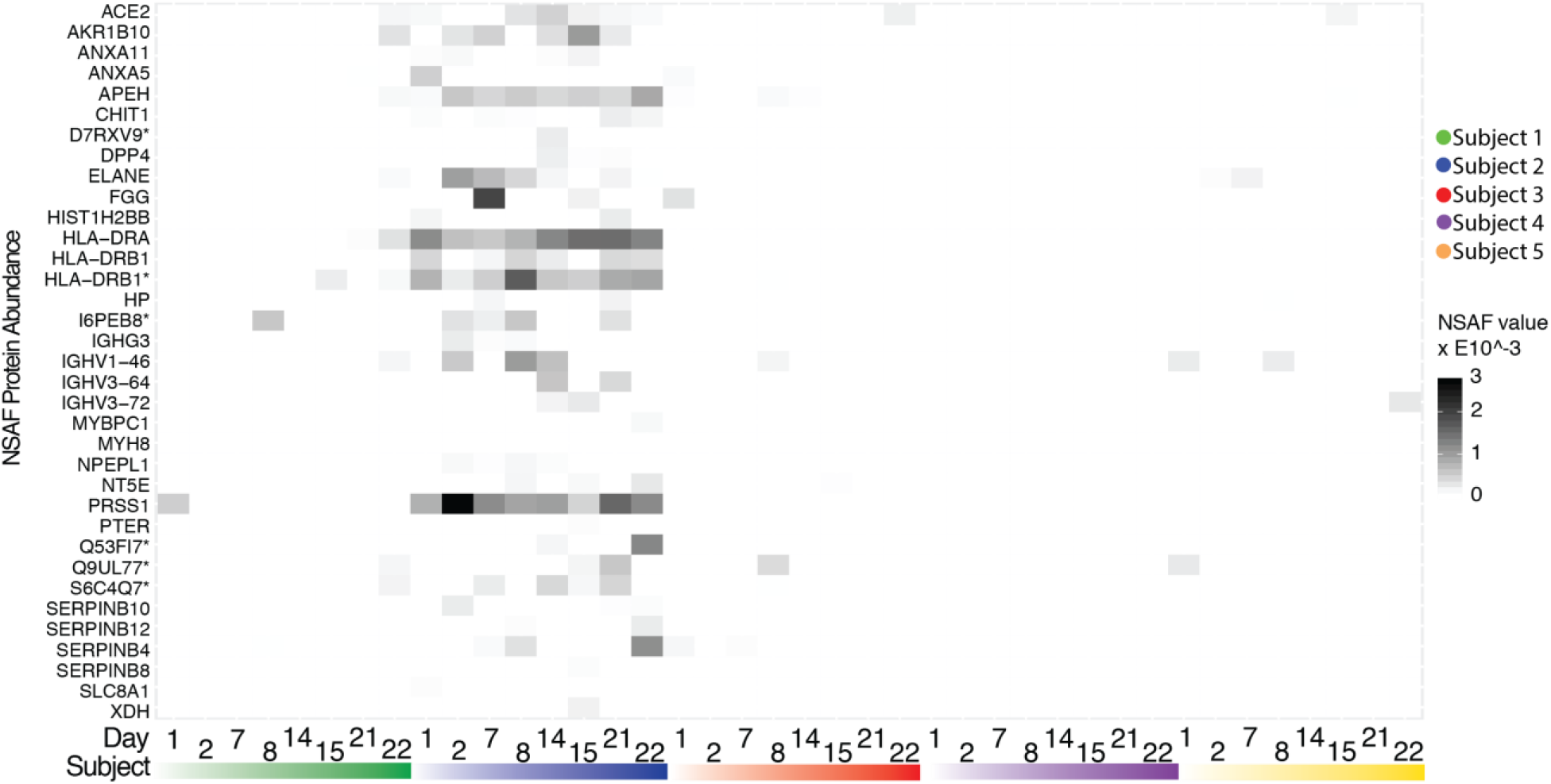
Upregulated inflammatory proteins greatest in Subject 2. Heatmap comparing protein abundances for subject 2 versus abundances in the other subjects. Proteins included in heatmap were in the top 50% summed Entropy for subject 2, and bottom 50% summed entropy (across each subject) for all other subjects. These proteins were inflammation-related like ELANE, MHC, and serine proteases.

## SUPPLEMENTAL TABLES

**Supplemental Table 1.**
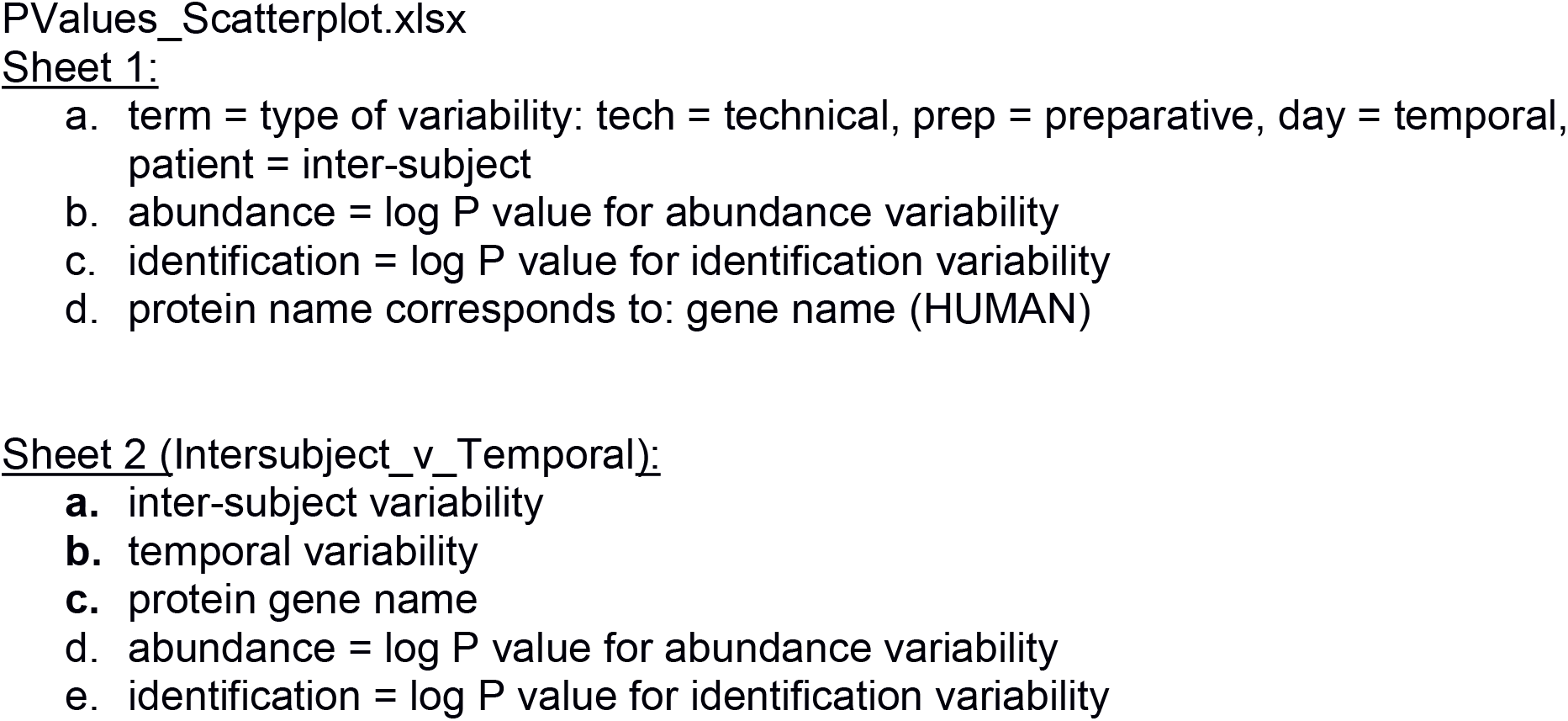
P values for each protein in the shared protein subset corresponding to each type of variability (technical, preparative, temporal, and inter-subject) and the respective abundance or identification consistency. On next sheet, table of significant inter-subject and temporal variability P values.

## Acknowledgements

We thank Parag Mallick for mass spectrometry advice. We thank members of the Elias laboratory for the insightful discussions and support, and especially the subjects of the study who made this research possible. This work was funded by the National Institutes of Health NIGMS Training Grant: 5T32GM008412 (EPC); Precision Health and Integrated Diagnostic Center at Stanford (JEE); National Institutes of Health: R01AI112401 (SH); National Science Foundation (NSF): DMS1501767; DHSC (SH); National Science Foundation Graduate Research Foundation: DGE-11474 (DS); National Institute of General Medical Sciences of the National Institutes of Health: T32GM007276 (DS); National Institute of Health:DP1OD000964,R01DE023113 (DAR).

## Materials and Methods

### Participants and Sampling Protocol

Healthy nonpregnant adults were recruited from the Stanford community, excluding individuals with chronic disease, hospitalization or antibiotic use in the previous 6 months, immunizations or international travel in the previous 4 weeks, or routine use of any prescription medication except birth control or hormone replacement therapy. These specimens were obtained with a protocol approved through the Stanford IRB: Protocol #25268. Characteristics of the five participants who completed the <u>sampling protocol</u> are summarized in <u>Table 1</u>. Participants collected ~2g stool specimens at home, which were frozen immediately without preservative in home freezers. Specimens were transferred without thawing to −80°C storage in the laboratory until further processing.

| Subject ID | Sex | Age | Racial Identification | BMI |
| --- | --- | --- | --- | --- |
| 1 | Male | 52 | Caucasian | 25.8 |
| 2 | Male | 45 | Caucasian | 27.5 |
| 3 | Male | 31 | Caucasian | 29.2 |
| 4 | Female | 31 | Caucasian | 20 |
| 5 | Female | 21 | Caucasian | 23.5 |

### Stool preparation for proteome analysis

Stool was thawed to aliquot 200mg per preparative replicate. Aliquots were solubilized in 1 mL 8M urea lysis buffer: 8M Urea (Invitrogen, CA; CAS# 57-13-6), 100mM NaCl(Fisher Scientific, CAS# 7467-14-5), 25mM Tris (Amresco, Solon, Ohio; CAS# 77-86-1), pH 8.2 with protease inhibitor cocktail (Roche Complete, EDTA free, Cat#05056489001. Pellets were physically disrupted by vortexing. Insoluble material was pelleted by a slow centrifugation step (2500 x g, 10 minutes). Microbes were pelleted by ultra centrifugation (35000 x g, 35 minutes). Resulting supernatant was then reduced with 5 mM DTT (20°C, 1 hour) and alkylated with iodoacetimide (15 mM, dark, 20°C, 1.5 hour). After quenching (5mM DTT, 20°C, 1 hour), trichloroacetic acid (TCA) was added to precipitate proteins (15% final volume, −20°C overnight). Acetone-washed protein pellets (3x) were resuspended in 8M Urea lysis buffer and separated by SDS-PAGE (1.5mm 4-12% Bis-Tris). Protein was allowed to migrate 0.5 cm into the gel before being stained with coomassie blue G250 (30 minutes), and de-stained overnight. De-stained gels were photographed and subject to densitometry-based quantification (Image J 1.49v). For in-gel proteolytic digestion, 5ng trypsin dissolved in 50mM AMBIC was added to diced and dehydrated gel pieces, and incubated overnight at 37°C. Digestion was quenched and extracted with 5% formic acid in 50% acetonitrile. Peptides were subsequently lyophilized by vacuum centrifugation, and de-salted using C-18 Sep-pak cartridges (Waters, Milford, CA; CAT# WAT036820). Peptides were resuspended in 1% formic acid, in volumes proportional to the densitometry measurements in which we compared the stool protein lane to the ladder lane and subsequently resuspended the peptides in the resulting ratio × 20uL. Peptides were then analyzed by LC-MS/MS as described below.

### Mass Spectrometry Analysis of the Human Gut Secreted Proteome

Peptides were resuspended in 1% formic acid. Resuspension volumes were determined for each specimen, based on Image J gel-lane normalization such that each specimen was estimated to have a concentration of 0.5ug/ul.

A Spark-Holland 920 autosampler loaded 1uL of resuspended peptides onto an 18 cm, 100 um, C18 fused micro-capillary column (inner diameter = 125 um) ending in an in-house pulled needle tip (internal diameter ~5 um). Ions were eluted from this column at 400 nL/min over a 172 minute gradient (4-25% 120 minutes, 25-45% 30 minutes, 45-95% 3 minutes, 98% 8 minutes, 95-3% 6.5 minutes) using an Agilent 1200 G1312 Binary pump (Agilent Technologies, San Jose, CA). The column was coupled to a home-built electrospray ionization source and LTQ-Velos Orbitrap mass spectrometer (Thermo Scientific, Santa Clara, CA. Full scans were collected in the orbitrap 60,000 resolution in centroid mode followed by a standard top-10, data-dependent framework with CID fragmentation in which singly charged precursor ions were excluded from fragmentation.. Dynamic exclusion was enabled and had a repeat count of 1 with a repeat duration of 30 seconds. All RAW data files were deposited in the PRIDE repository (30) and assigned the accession ID PXD010491 (Username; Password: vlXOIpLT)

### Protein FDR Filtering and normalization

All human proteins and common contaminants were downloaded from Uniprot ((31)) and concatenated with a standard peptide list totaling to a database with 423,268 entires Uniprot, downloaded December 1, 2014. The msConvert program was used to generate peak lists from the original data. All protein sequences were reversed and concatenated to the original forward-oriented proteins ((32)). Spectra were assigned to peptides using a semispecific enzyme specificity with the SEQUEST search algorithm (33) using static carbamidomethylation of cysteines, differential oxidized methionines, 50 ppm precursor masstolerance, 2 miscleavages and .0.8 Da fragment ion tolerance. FDRs were estimated with the target-decoy search strategy and peptide false-positives were filtered to 1% and protein FDR was filtered to (1.5%). Protein spectral counts were normalized according to NSAF (34).

### Variability Significance Calculations

Variability testing was performed on the shared protein subset of 74 proteins with R version 3.5.1 and associated packages as noted below. These proteins were evaluated with two types of significance testing: ANOVA (continuous non-zero values) and general linearized model (presence vs absence). First, each raw mass spectrometry output data file was assigned a unique combination of experimental identifiers: technical replicate number; preparative replicate number; time point; and subject number. Proteins identified from each raw file were assigned the corresponding identifier attributes, along with the protein’s log(NSAF) abundance.

To calculate abundance-based p values with ANOVA, we only considered non-zero abundance values for each protein, and for each specific attribute being tested. For example, when testing technical replicate variability, only pairs of non-zero abundance values would be considered; no technical replicates would be considered for preparative replicate 1 of time point 2, subject 3 if only one technical replicate was recorded with a non-zero abundance. Linear models were generated with the “lm” function (package: stats version 3.5.1) considering all valid response variables as described above, at the technical, preparative, time point, and subject levels. ANOVA p values were calculated from the resulting linear models with the “anova” function (package: stats version 3.5.1).

We considered the entire data matrix to calculate binary identification p values with a logistic regression (glm). All non-zero abundance values were thresholded as present (1); all remaining zero values were considered absent (0). Logisitic regression p values were calculated from all response variables thresholded as above, with the “glm” function (package: stats version 3.5.1). P values from both variance tests were compared using ggplot version 3.0.0).

### Microbiome Analysis

Stool specimens were collected at home by participants and frozen immediately in home freezers (~−20°C), transferred frozen to the laboratory, then stored at −80°C until further processing. DNA was extracted from ~200 mg stool using the QIAGEN AllPrep DNA/RNA 96 Kit (Cat # 80311) following the manufactures instructions, with the addition of a 60 second bead-beating step using the FastPrep-24 5G Benchtop Homogenizer at speed 6.5. The V4-V5 region of the 16S rRNA gene was amplified using a forward primer 515FB (5′-GTGYCAGCMGCCGCGGTAA-3′) with an error-correcting barcode and 926R (5′-CCGYCAATTYMTTTRAGTTT-3′). PCR products were then purified, pooled in equimolar concentrations, and sequenced on two 2 × 250 Illumina HiSeq paired-end runs.

After demultiplexing, raw reads were quality trimmed using the DADA2 pipeline (DADA2 version 1.3.4) in R (R version 3.2.4). Forward reads were truncated to 240 bp, and reverse reads to 220 bp, and quality filtered with following settings: maxN=0, maxEE=2, truncQ=2. Following quality trimming, Amplicon Sequence Variants (ASVs) were inferred using the DADA2 pipeline, forward. Forward and reverse reads were then merged, and chimeras were removed using DADA2’s consensus option. Both ASV tables from different sequencing runs were then merged, and ASV sequences shorter than 353 bp or longer than 368 bp were discarded.

Taxonomy was assigned to each ASV using the RDP classifier and the RDP Trainset version 14. ASV sequences were then aligned using ClustalW as implemented in the msa R package (msa version 1.6.0), from which a phylogenetic tree was inferred by the phangorn R package (phangorn version 2.2.0). The ASV tablecounts, subject dataand timepoint identifiers, taxonomy assignments, phylogenetic tree, and ASV sequences were then bundled into a single phyloseq data object for further plotting and statistical analysis (phyloseq version 1.19.1).

### Principal Coordinate and Component Analysis

Stool proteins and microbes were plotted on separate ordination graphs to examine inter-individual and temporal trends in both the proteome and microbiome. For the proteins, technical and preparative replicates were averaged to consider only the temporal and inter-subject specimens: each of the five subjects had eight stool specimens for each time point. All proteins were used to produce all protein ordination graphs. 1213 microbes met or surpassed the following criteria: 5% presence and 10 count abundance and were subsequently used to produce all microbe ordination graphs.

PCoA were generated using the R package phyloseq version 1.24.2. Visualization was created using ggplot2 version 3.0.0. Bray Curtis and Euclidean distances were used to assess protein separation. Bray Curtis, Unifrac, and unweighted Unifrac were used to assess microbe separation.

Temporal patterns within protein and microbe data sets were created based on the centered (mean set to 0) Bray Curtis PCoA vectors from the Bray Curtis PCoA ordination plots, respectively. Time series are chronologically represented as a transition from white to black. No trends were apparent from either data source. Euclidean distances between each time point were summed to quantify the extent to which subjects’ specimens were temporally divergent.

